# A bicistronic *Aldh1a3-P2A-TagBFP* knock-in reporter mouse line for studying genitourinary tract development

**DOI:** 10.64898/2026.09.21.752752

**Authors:** Aurore Chemla, Annie Rubio-Ortega, Lauren Brûlé, Eduardo Carmona-Miranda, Mia Hudon, Maxime Bouchard, Oraly Sanchez-Ferras

## Abstract

Aldehyde dehydrogenase 1a3 (Aldh1a3) is an enzyme involved in retinoic acid synthesis with dynamic expression patterns during development, including in the urogenital system. Here, we generated a bicistronic *Aldh1a3-P2A-TagBFP* knock-in mouse using CRISPR/Cas9 genome editing, inserting *TagBFP* immediately upstream of the endogenous *Aldh1a3* stop codon. Correct targeting was confirmed by Oxford Nanopore long-read sequencing, and heterozygous and homozygous mice were viable and fertile without overt morphological abnormalities. TagBFP fluorescence faithfully overlapped with endogenous Aldh1a3 immunoreactivity and reproduced established expression domains in the developing craniofacial region, intestine, kidney, and broader urogenital system. Extensive characterization of the urogenital system revealed dynamic, spatially restricted BFP reporter activity in *Aldh1a3*-expressing domains across several key structures, including the ureteric bud and collecting duct lineage, seminal vesicles, caput epididymis, and developing uterine horns. The *Aldh1a3-P2A-TagBFP* mouse provides a fluorescent resource for visualizing *Aldh1a3* expression across development and in adult tissues, including for the characterization of *Aldh1a3*-expressing domains in the urogenital system. The relatively low fluorescence intensity of TagBFP should be considered when assessing low-level reporter expression.

## Introduction

Aldehyde dehydrogenase 1a3 (Aldh1a3; also known as Raldh3) is a key enzyme in retinoic acid (RA) biosynthesis that catalyzes the irreversible oxidation of retinaldehyde to RA, driving essential developmental transcriptional programs and cellular differentiation^1,2^. Loss of *Aldh1a3* in mice results in neonatal lethality with severe craniofacial and sensory malformations^3^, while combined inactivation with *Aldh1a2* severely disrupts craniofacial, central nervous system, and kidney development, beyond the defects seen with *Aldh1a2* deficiency alone^4–6^. These findings demonstrate that Aldh1a3 plays a critical role in driving RA-dependent organogenesis.

While the related enzymes Aldh1a1 and Aldh1a2 display broad epithelial and mesenchymal distribution, respectively^1,4,7^, Aldh1a3 exhibits a distinct, spatially restricted expression profile in specific embryonic structures, including regions of the craniofacial system, intestine, eye, and urogenital tract^1,4,8^. Within the developing urogenital system, *Aldh1a3* is expressed in the nephric duct (ND), ureteric bud derivatives, and seminal vesicle epithelium^1,4,9^. Our previous single-cell transcriptomic profiling identified *Aldh1a3* as a defining marker of a distinct, ND progenitor population (NdPr4) located at the leading edge of the ND that subsequently contributes to ureteric bud initiation^9^. These observations indicate that *Aldh1a3* expression is dynamically regulated during urogenital development. However, the spatial distribution and temporal dynamics of *Aldh1a3* expression during organogenesis remain incompletely characterized, partly because of the limited availability of tools for directly visualizing *Aldh1a3*-expressing cells in developing tissues.

Existing genetic approaches for studying *Aldh1a3*-expressing populations include inducible CreERT2 knock-in alleles and lacZ-based reporter lines^10,11^. These models have provided valuable insights into *Aldh1a3* expression and, in the case of Cre-based approaches, lineage contribution. CreERT2-based strategies require breeding with Cre-dependent reporter strains and temporal induction by tamoxifen, while lacZ-based reporters have been particularly useful for histological visualization of *Aldh1a3* expression in fixed tissues. A fluorescent reporter expressed directly from the endogenous *Aldh1a3* locus therefore provides a complementary approach for visualizing *Aldh1a3*-expressing cells and their distribution during development using fluorescence microscopy.

Here, we generated a bicistronic *Aldh1a3-P2A-TagBFP* knock-in mouse line by inserting a *P2A-TagBFP* cassette immediately upstream of the endogenous *Aldh1a3* stop codon. The *P2A* sequence promotes ribosomal skipping during translation, enabling the production of separate Aldh1a3 and TagBFP proteins from a single *Aldh1a3*-derived transcript. This strategy places TagBFP under the control of endogenous *Aldh1a3* regulatory elements while maintaining the endogenous *Aldh1a3* coding region upstream of the P2A-TagBFP cassette. We validated the targeted allele by Oxford Nanopore long-read sequencing and assessed the viability and gross morphology of heterozygous and homozygous animals. We further characterized TagBFP expression across multiple developmental stages of the kidney and reproductive tract and evaluated its overlap with endogenous Aldh1a3 protein expression. This reporter line provides a fluorescent resource for investigating the spatial and temporal distribution of *Aldh1a3*-expressing cells during urogenital development, including the developing kidney, while also enabling the study of *Aldh1a3* expression in other embryonic and adult tissues where the gene is expressed.

## Results

### Generation and initial characterization of a bicistronic *Aldh1a3* reporter mouse line

To enable direct visualization of *Aldh1a3*-expressing cells, we generated a bicistronic *Aldh1a3-P2A-TagBFP* knock-in mouse line using CRISPR/Cas9-mediated genome editing. A *P2A-TagBFP* cassette was inserted immediately upstream of the endogenous *Aldh1a3* stop codon in exon 13, placing *TagBFP* expression under the control of the endogenous *Aldh1a3* regulatory elements while preserving the *Aldh1a3* coding sequence (**Figure 1A**). The *P2A* sequence promotes ribosomal skipping during translation, enabling the production of separate Aldh1a3 and TagBFP proteins from the endogenous *Aldh1a3* transcriptional unit. To facilitate precise integration at the intended site, a guide RNA targeting *Aldh1a3* close to the endogenous stop codon was designed using CRISPOR. Guide selection considered predicted on-target activity and potential off-target sites. Guide activity was first evaluated by assessing target-site modification in transient blastocysts generated following CRISPR/Cas9-mediated editing. Genomic PCR, Sanger sequencing, and Tracking of Indels by Decomposition (TIDE) analysis were used to assess editing efficiency at the target site (**Figure S1**).

**Figure 1.**
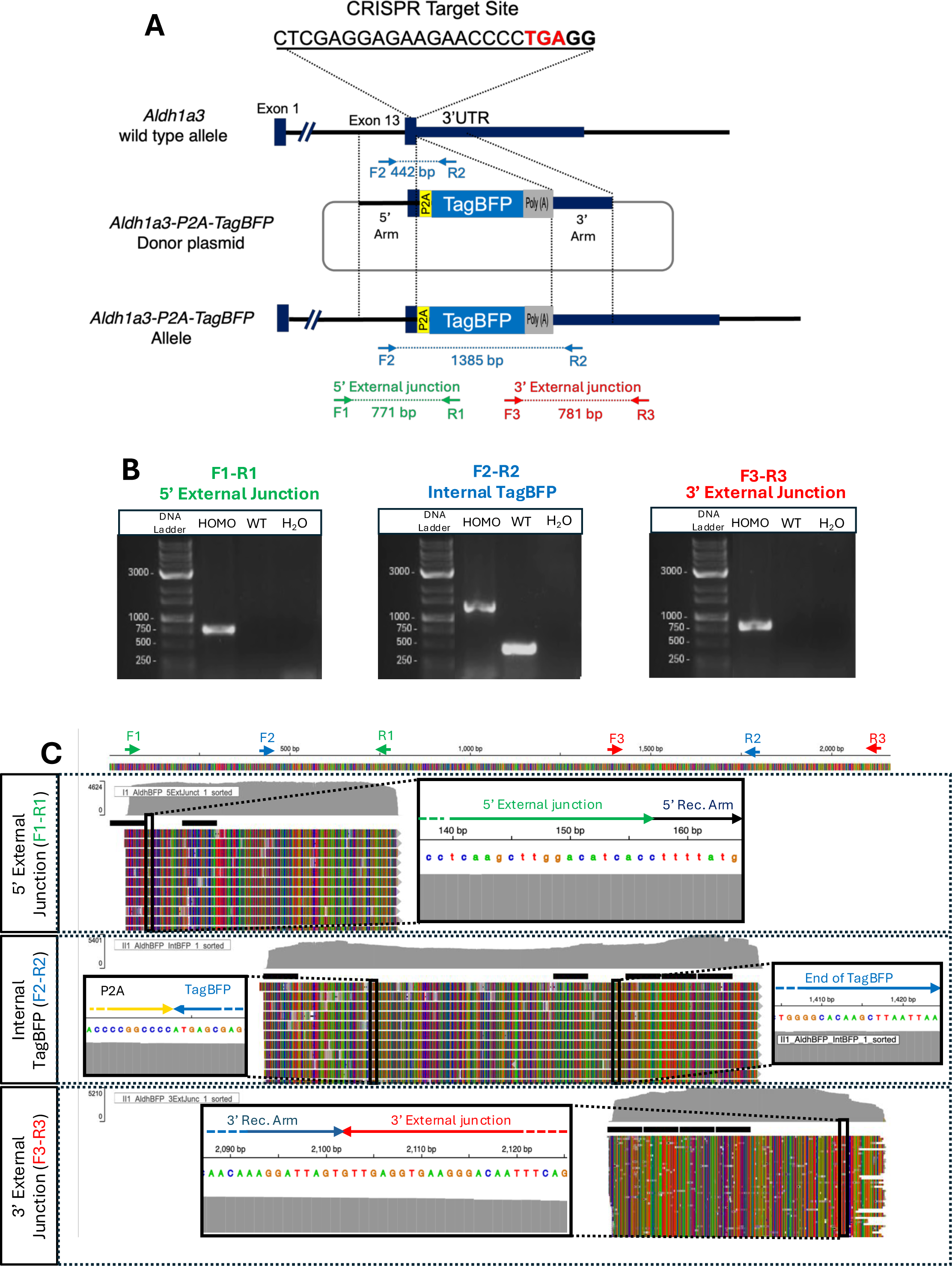
Generation and validation of the *Aldh1a3-P2A-TagBFP* knock-in mouse model. **A.** CRISPR/Cas9 targeting strategy. The guide RNA target site was designed in *Aldh1a3* exon 13, spanning the endogenous stop codon (red bold) and adjacent *PAM* sequence (blue). The donor plasmid contained a 5′ homology arm, *P2A* sequence (yellow), *TagBFP* coding sequence (blue), *SV40 poly(A)* signal (gray), and 3′ homology arm. Homology-directed repair (HDR) resulted in insertion of the *P2A-TagBFP-poly(A)* cassette immediately upstream of the endogenous stop codon, enabling bicistronic expression of *Aldh1a3* and *TagBFP* from the endogenous *Aldh1a3* locus. Primer pairs used for genotyping of the 5′ external junction (F1–R1, green), internal *TagBFP* region (F2–R2, blue), and 3′ external junction (F3–R3, red) are indicated. **B.** PCR validation of the targeted allele. Genomic DNA from *Aldh1a3-P2A-TagBFP* homozygous mice (HOMO), wild-type (WT) mice, and no-template control (H₂O) was amplified using the indicated primer pairs. F1–R1 and F3–R3 specifically detected the targeted allele, while F2–R2 amplified a 442 bp wild-type allele and a 1385 bp knock-in allele. **C.** Oxford Nanopore long-read sequencing validation of the *Aldh1a3-P2A-TagBFP* knock-in allele. Representative alignments across the targeted locus confirm precise integration of the *P2A-TagBFP* cassette and correct HDR-mediated recombination. Sequencing reads spanning the 5′ external junction (top), internal *P2A-TagBFP* region (middle), and 3′ external junction (bottom) demonstrate seamless integration and structural integrity of the targeted allele. Insets show nucleotide sequences across the homology arm recombination sites. Grey histograms indicate read coverage, and colored reads represent individual Oxford Nanopore reads aligned to the reference knock-in allele.

The donor construct containing the 5′ and 3′ homology arms, the *P2A-TagBFP* cassette, and an *SV40* polyadenylation signal was introduced together with Cas9 and the guide RNA into two-cell-stage C57BL/6NCrl embryos. Seven founder animals were screened by PCR using primers targeting internal and external regions of the integration site (**Figure 1A**; **Table S1**). Three founders displayed the expected junction PCR products, consistent with targeted integration. One founder was confirmed to transmit the knock-in allele through the germline and was subsequently selected for colony establishment and further characterization.

To evaluate germline transmission, offspring from heterozygous × wild-type crosses were genotyped. Among 60 progeny analyzed, 36 were wild-type (60%) and 24 were heterozygous (40%). This distribution did not significantly deviate from the expected 1:1 Mendelian ratio (χ² = 2.4, p = 0.12; **Table 1**). The sex distribution was balanced, with 28 males (46.7%) and 32 females (53.3%) observed among the offspring. F1 heterozygous mice were subsequently intercrossed to generate F2 progeny, including wild-type, heterozygous, and homozygous animals (n = 10; **Table 1**).

**Table 1.**
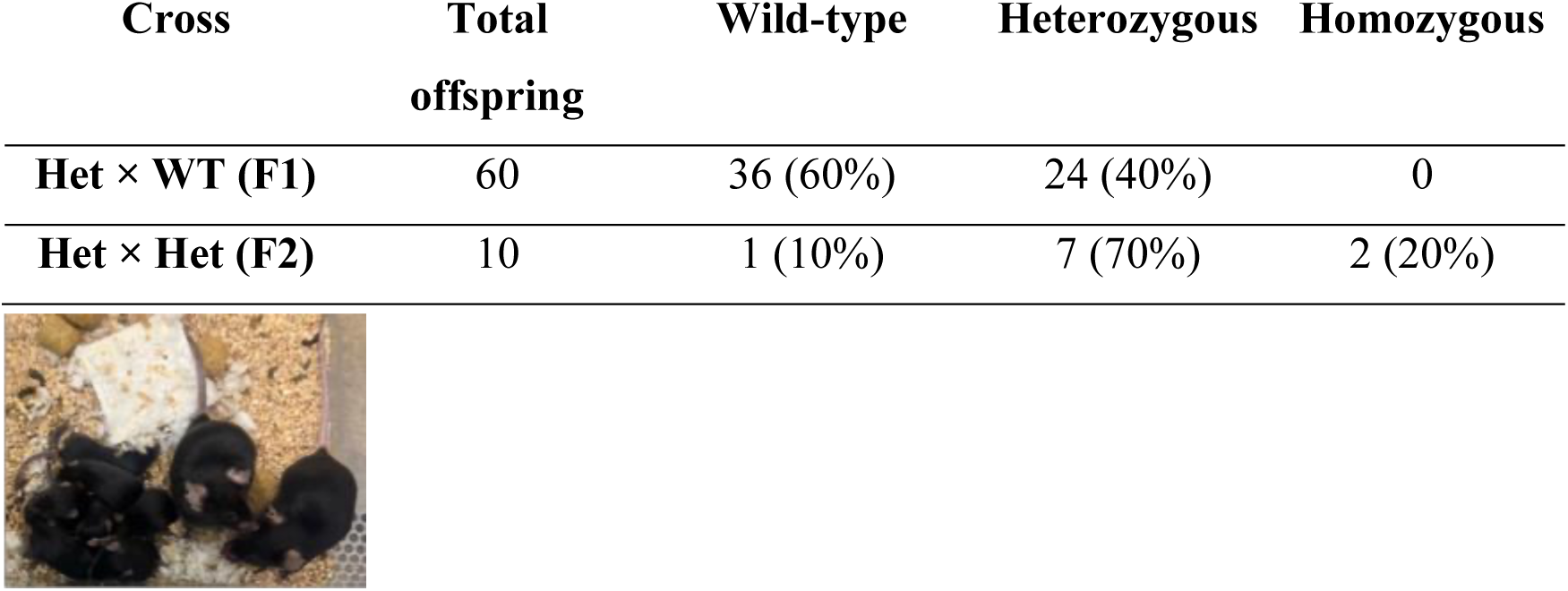
Genotypic distribution of offspring from *Aldh1a3-P2A-TagBFP* reporter mouse crosses and representative picture of homozygous breeding pair with offspring.

The homozygous knock-in allele was further evaluated by PCR and Oxford Nanopore long-read sequencing. These analyses confirmed the expected integration of the *P2A-TagBFP-ploy(A)* cassette and the integrity of the analyzed 5′ and 3′ junctions’ regions at the targeted locus (**Figure 1B, C; Figure S2**). Homozygous animals were subsequently used to establish the reporter colony. During colony establishment, homozygous breeding pairs produced 144 viable offspring. Under the conditions examined, heterozygous and homozygous reporter mice were viable and capable of reproduction, with no obvious abnormalities in growth or gross adult morphology (**Table 1**).

TagBFP fluorescence was detected in progeny derived from the germline-transmitting founder and was observed in expression domains consistent with previously described *Aldh1a3* expression patterns at E9.5, including the rostral cranial region and the nephric duct (ND) ^9,12^ (**Figure 2A**). This pattern is consistent with our previous identification of *Aldh1a3* as a marker of the caudal NdPr4 leader-cell population during ND collective migration^9^. Immunostaining for TagBFP and Aldh1a3 showed overlapping signal in both the cranial and caudal ND regions, supporting the correspondence between reporter fluorescence and endogenous Aldh1a3 protein expression (**Figure 2B**; **Figure S3**).

**Figure 2.**
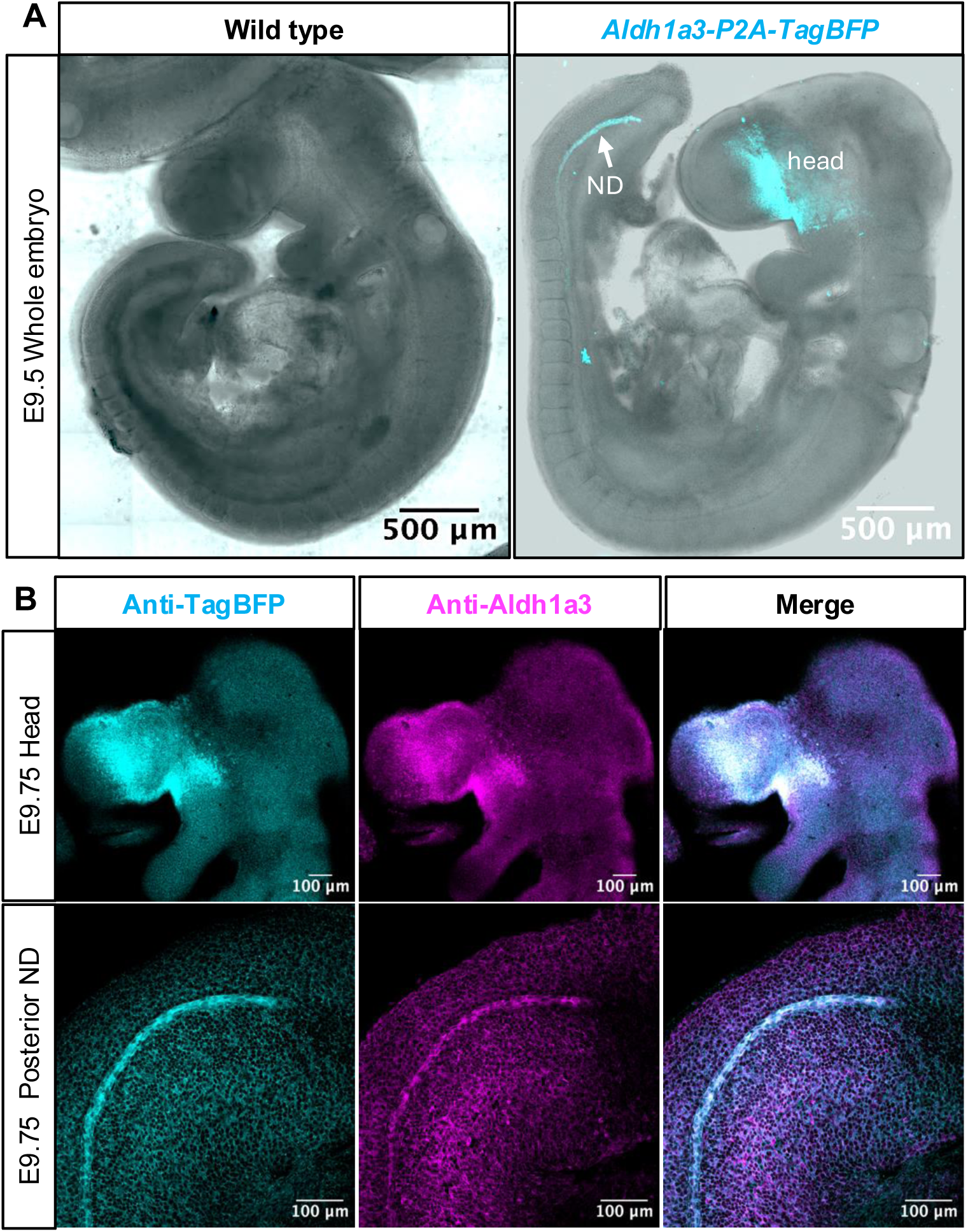
Characterization and validation of *Aldh1a3-P2A-TagBFP* expression in E9.5 mouse embryos. **(A)** Whole-mount fluorescence confocal imaging of E9.5 wild type (left) and *Aldh1a3-P2A-TagBFP* (right) mouse embryos. TagBFP signal (cyan) is specifically localized to the caudal nephric duct (ND) and the rostral head region, recapitulating endogenous *Aldh1a3* expression. **(B)** Co-immunostaining with anti-TagBFP (cyan) and anti-Aldh1a3 (magenta) antibodies in E9.75 whole-mount embryos demonstrates co-localization of the TagBFP reporter with endogenous Aldh1a3 protein in both the head and caudal ND domains. Scale bars: 200 µm (whole embryos), 500 µm (E9.5 brightfield overlay), and 100 µm (immunostaining panels).

We next assessed TagBFP expression in tissues previously reported to express *Aldh1a3* during embryonic and postnatal development^1,12,13^. In the craniofacial region, TagBFP fluorescence was detected around the developing eye and facial ectoderm at E9.75 and became progressively restricted to defined ocular structures between E10.0 and E12.5 (**Figure 3A–D**)^12^. Reporter fluorescence was also detected in the epithelium of the E18.5 small intestine and in the kidneys and seminal vesicles of the P5 male urogenital system^1,4^ (**Figure 3 E,F**). As an additional control, P5 wild type kidneys imaged using the same acquisition parameters showed no detectable TagBFP fluorescence (**Figure S4**).

**Figure 3.**
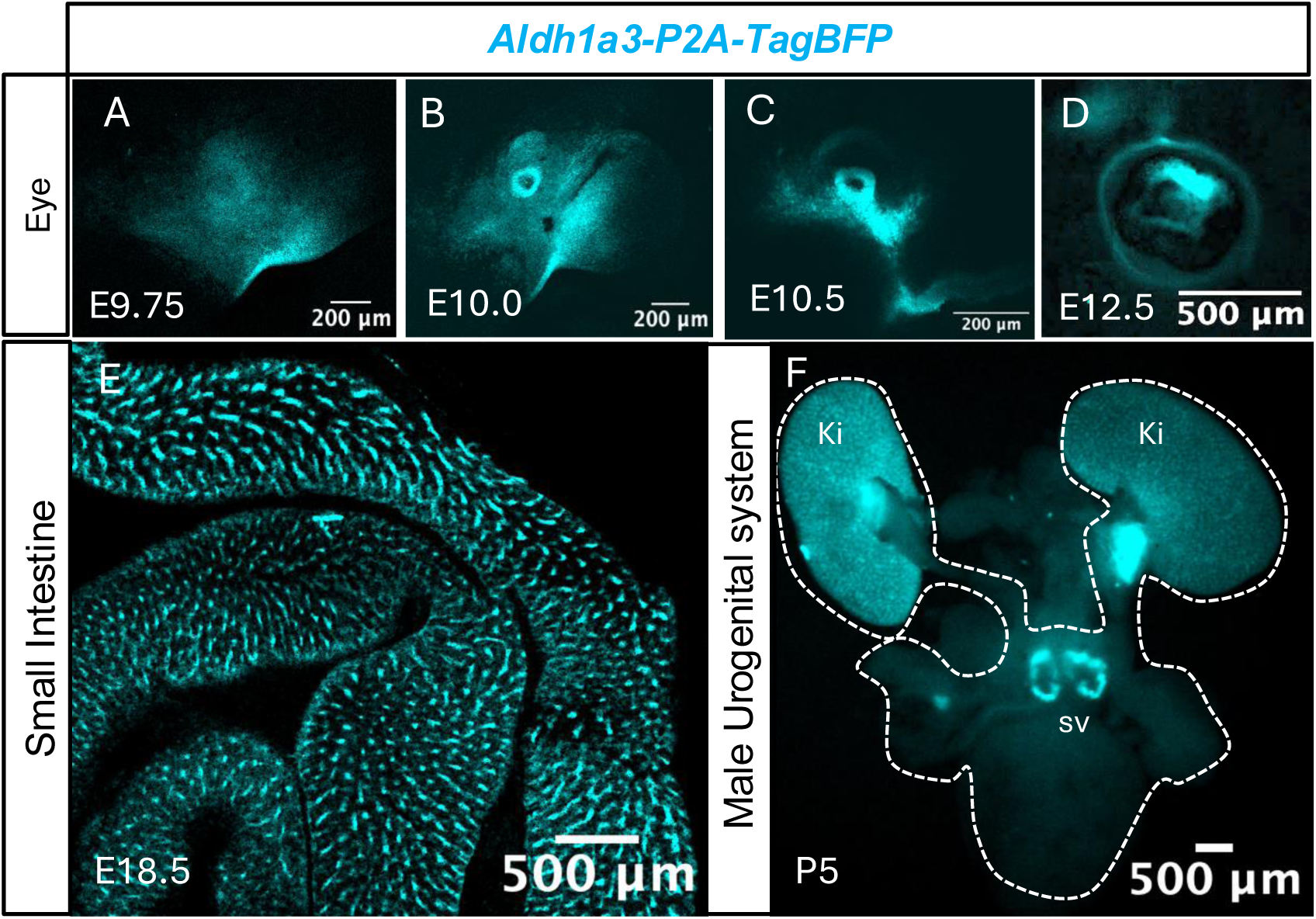
Fluorescence reporter expression of the *Aldh1a3-P2A-TagBFP* reporter line across different developing organs. Whole-mount TagBFP fluorescence imaging of *Aldh1a3-P2A-TagBFP* mouse tissues. **(A–D)** Spatial expression dynamics during optic cup and eye development at E9.75 (**A**), E10.0 (**B**), E10.5 (**C**), and E12.5 (**D**). **(E)** Robust reporter fluorescence outlining intestinal villi in the E18.5 small intestine. **(F)** Expression in the P5 male urogenital system (outlined by dashed white line), showing reporter signal in the kidneys (Ki), seminal vesicles (sv). Scale bars: 200 µm (A–C), 500 µm (D–F).

Together, these results demonstrate successful germline transmission and establishment of a viable *Aldh1a3-P2A-TagBFP* reporter line that recapitulates previously described *Aldh1a3* expression domains and enables direct visualization of *Aldh1a3*-expressing tissues.

### Fluorescence reporter expression during kidney development

*Aldh1a3* is expressed during kidney development, where it contributes to RA signaling through RA synthesis^1,4,14^. We therefore examined the spatial distribution of TagBFP fluorescence from the *Aldh1a3-P2A-TagBFP* reporter in the developing mesonephros and metanephros across embryonic and postnatal stages.

During mesonephros development, TagBFP fluorescence was restricted to the posterior region of the nephric duct (ND) at E9.5 and E10.5, highlighting the site of ureteric bud emergence (**Figure 4A–B**). By E11.5, reporter expression extended further along the anterior ND axis, establishing a clear posterior-to-anterior gradient (**Figure 4C–D**). High signal intensity was concentrated in the caudal ND, ureter, and ureteric bud (UB), while expression progressively diminished toward the more anterior regions of the ND (**Figure 4C–D**).

**Figure 4.**
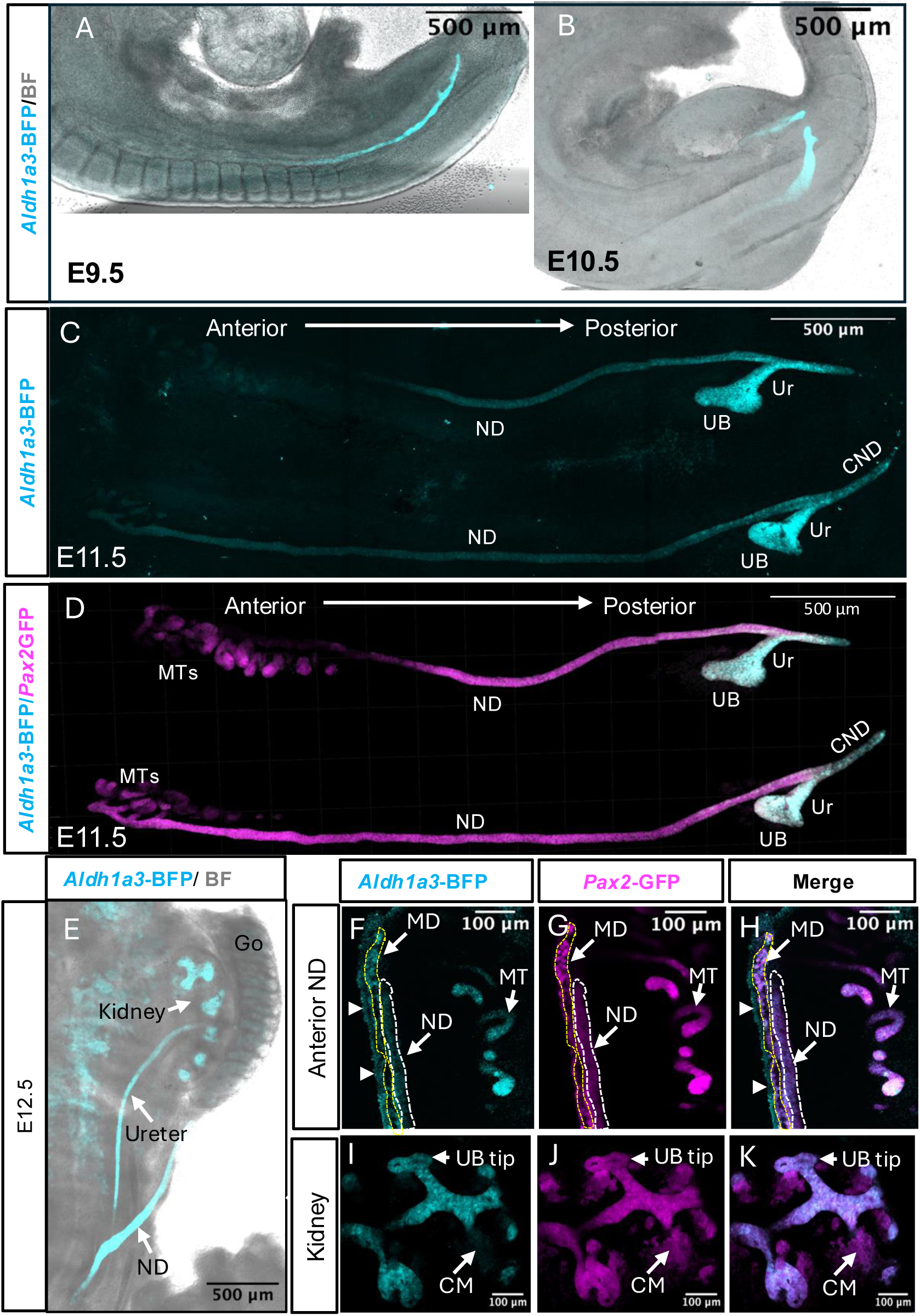
*Aldh1a3-P2A-TagBFP* reporter expression during mesonephros and metanephros development. (**A–E**) Whole-mount confocal images showing the spatial distribution of *Aldh1a3*-TagBFP across developmental stages: E9.5 (**A**), E10.5 (**B**), E11.5 (**C, D**), and E12.5 (**E**). Panels A, B, and E show TagBFP signal overlaid with brightfield (BF). Panel D shows a 3D reconstruction of an E11.5 dual-reporter urogenital system (UGS) displaying *Aldh1a3*-TagBFP (cyan) and *Pax2*-GFP (magenta). (**F–K**) High-magnification confocal images of E12.5 *Aldh1a3-P2A-TagBFP*; *Pax2-GFP* dual-reporter embryos displaying TagBFP signal (cyan; F, I), Pax2-GFP signal (magenta; G, J), and channel overlays (H, K). (**F–H**) Anterior mesonephric region highlighting mesonephric tubules (MT), nephric duct (ND; white dashed outline), and Müllerian duct (MD; yellow dashed outline and arrowheads). (**I–K**) Developing metanephric kidney highlighting ureteric bud tips (UB tip) and cap mesenchyme (CM). Abbreviations: Go: gonad; UB: ureteric bud; Ur: ureter; ND: nephric duct; CND: common nephric duct; MD: Müllerian duct; MT/MTs: mesonephric tubules; CM: cap mesenchyme. Scale bars: 500 µm (A–C, E), 100 µm (D, F–K).

At E12.5, TagBFP expression was maintained along the entire ND axis as well as its mesonephric and metanephric derivatives (**Figure 4E–K**). High-magnification analysis of the anterior ND region (Figure 4F-H) revealed TagBFP fluorescence extending along the anterior ND (white dashed outline) and within the mesonephric tubules. Reporter signal was also prominently detected in the developing Müllerian duct epithelium (MD; yellow dashed outline), where it co-localized with *Pax2-*GFP, as well as in the adjacent *Pax2*-GFP-negative periductal mesenchyme (indicated by arrowheads in **Figure 4F-G**). In the caudal metanephric region, whole-mount imaging (**Figure 4E**) demonstrated prominent TagBFP signal along the ureter and caudal ND, while fluorescence remained low or absent in the developing gonads. Furthermore, in the E12.5 metanephric kidney, TagBFP fluorescence was specifically restricted to the branching ureteric bud tips (**Figure 4I-K**), whereas signal was absent in the adjacent *Pax2*-GFP-positive cap mesenchyme (**Figure 4I-K**). During metanephric kidney development, *Aldh1a3*-TagBFP fluorescence reveals a complex, branching tubular distribution characteristic of the UB-derived collecting duct lineage (**Figure 5A–D**). Reporter fluorescence extended from the central collecting system in the renal pelvis region throughout the collecting duct network **(Figure 5E–G)**. The signal was not uniform throughout the collecting ducts system and appeared more prominent in distal cortical collecting ducts near the kidney periphery (**Figure 5 E-G**). Co-localization with *Pax2*-GFP further supported the localization of TagBFP fluorescence within the broader collecting duct lineage (**Figure 5E– G**). In high-magnification views of the P5 renal cortex, *Pax2*-GFP labeled both the collecting duct network and adjacent nephron structures, whereas TagBFP fluorescence was primarily associated with the branching collecting duct tree and was not readily detected in neighboring Pax2-positive nephron structures (**Figure 5H-J, white arrowhead**).

**Figure 5.**
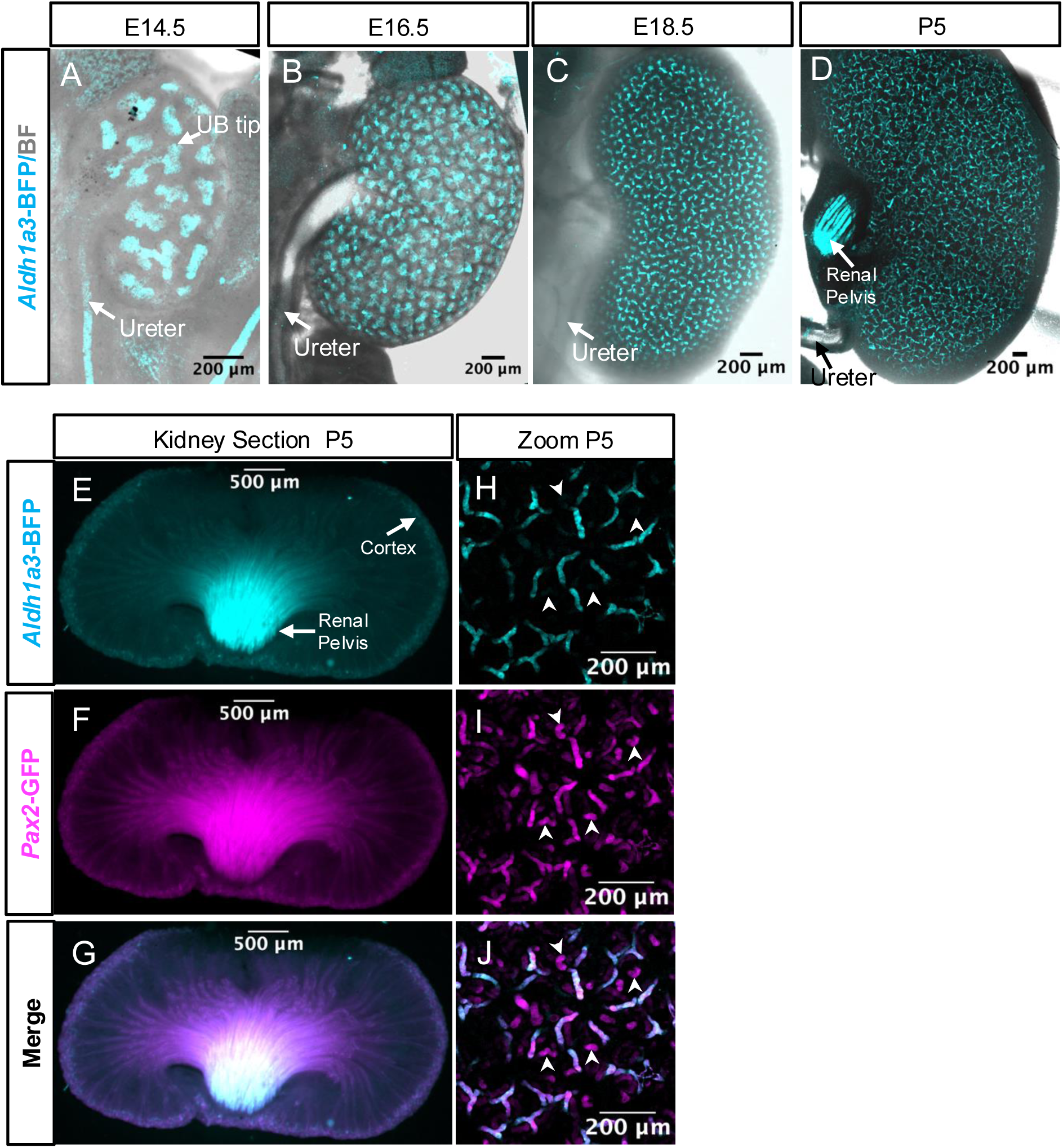
Spatiotemporal *Aldh1a3-P2A-TagBFP* reporter expression during metanephric kidney development. (**A–D**) Whole-mount fluorescence confocal images (overlayed with brightfield) showing *Aldh1a3*-TagBFP expression (cyan) in embryonic and postnatal kidneys at E14.5 (**A**), E16.5 (**B**), E18.5 (**C**), and P5 (**D**). Labels indicate the ureteric bud tips (UB tip) and ureter. (**E–G**) Stereomicroscope fluorescence images of P5 dual-reporter kidney vibratome sections displaying *Aldh1a3*-TagBFP signal (cyan; **E**), *Pax2*-GFP signal (magenta; **F**), and channel overlay (**G**). Anatomical labels indicate the renal cortex and renal pelvis. (**H–J**) High-magnification confocal views of the P5 renal cortex showing *Aldh1a3*-TagBFP signal (cyan; **H**), *Pax2*-GFP signal (magenta; **I)**, and channel overlay (**J**). White arrowheads indicate *Pax2*-GFP– positive renal tubules lacking *Aldh1a3*-TagBFP reporter signal. Scale bars: 200 µm (**A–D**, **H–J**), 500 µm (**E–G**).

Reporter fluorescence in the ureters was strong from E11.5 through E14.5 and appeared reduced at later stages, including E16.5, consistent with previous observations^14^ (**Figure 4C-E and 5A-D, Figure S5**). The distribution of Aldh1a3-TagBFP fluorescence was comparable between male and female embryos and between left and right kidneys, with no obvious differences detected under the conditions examined (**Figure S5**).

Together, these observations show that *Aldh1a3*-TagBFP fluorescence is detected in several epithelial components of the developing mesonephros and metanephros, including the ureteric bud, collecting duct network, ureter, nephric duct, mesonephric tubules, and Müllerian duct. The reporter signal was more prominent in selected regions of the collecting duct system and was not readily detected in the cap mesenchyme or adjacent nephron structures under the imaging conditions used.

### Spatiotemporal dynamic of *Aldh1a3*-TagBFP fluorescence in the reproductive tract

RA signaling contributes to the development of the reproductive tract and to the morphogenesis of urogenital structures^2,4,15–18^. To characterize the distribution of Aldh1a3 expression during reproductive tract development, we examined TagBFP fluorescence from the *Aldh1a3-P2A-TagBFP* reporter in male and female urogenital tissues across embryonic, postnatal, and adult stages.

*Male reproductive tract*. At the early embryonic stage examined (E12.5), TagBFP fluorescence was detected along the ND and in the developing rete testis located at the gonad–mesonephros interface (**Figure 6A,A′**). Reporter fluorescence was not readily detected in the gonadal parenchyma at this stage.

**Figure 6.**
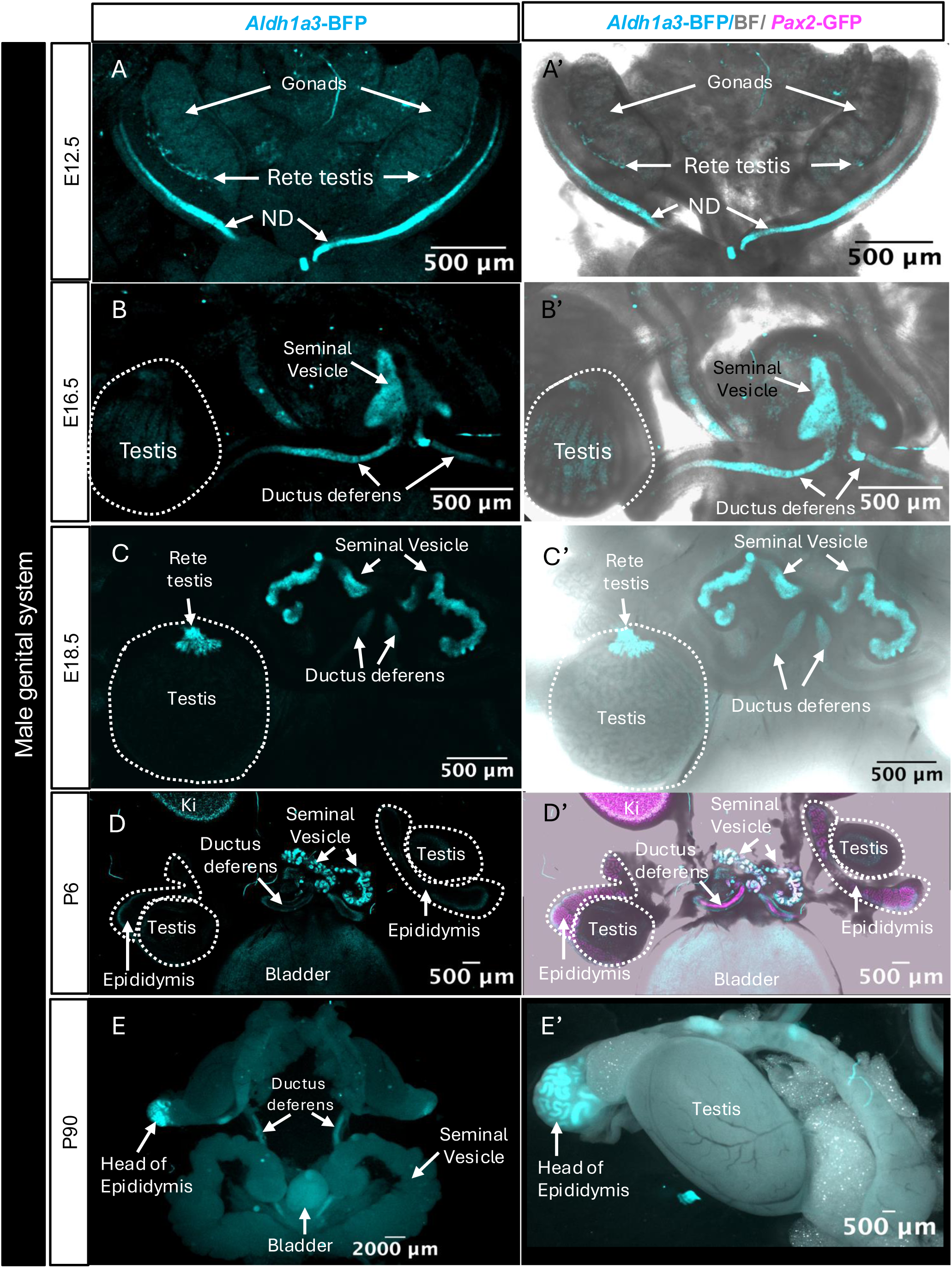
*Aldh1a3-P2A-TagBFP* reporter expression during male genital tract development. (**A–E′**) Whole-mount fluorescence confocal and brightfield overlay images showing *Aldh1a3*-TagBFP expression (cyan) alongside *Pax2*-GFP (magenta; **D′**) across embryonic, postnatal, and adult stages of male genital tract development. Left panels ( **A–E**) display *Aldh1a3*-TagBFP fluorescence; right panels (**A′–E′**) show corresponding channel overlays. (**A, A′**) E12.5 embryonic reproductive tract displaying labels for the gonads, rete testis, and nephric duct (ND). ( **B, B′**) E16.5 male genital tract with labels indicating the testis (dashed outline), seminal vesicle, and ductus deferens. (**C, C′**) E18.5 male reproductive structures highlighting the rete testis, seminal vesicles, and ductus deferens. (**D, D′**) Postnatal day 6 (P6) male urogenital region showing *Aldh1a3*-TagBFP (cyan) and *Pax2*-GFP (magenta; **D′**) with labels for the kidney (Ki), epididymis, testis (dashed outline), seminal vesicle, ductus deferens, and bladder. ( **E, E′**) Adult male genital tract whole-mount view (**E**) and high-magnification view (**E′)** displaying the head of the epididymis, ductus deferens, testis, and bladder. ND: Nephric duct; Ki: Kidney. Scale bars: 500 µm (A–D′, E′), 2000 µm (E).

As male reproductive tract development progressed, TagBFP fluorescence was detected in the budding lumen of the seminal vesicles, the rete testis, and the ductus deferens (**Figure 6B–C′**). Further comparison of *Aldh1a3-P2A-TagBFP* and *Pax2-GFP* reporter expression at Postnatal day (P)6 showed distinct patterns along ND-derived structures (**Figure 6D, D′**). Whereas *Pax2*-GFP broadly labeled the epithelium of the ductus deferens, seminal vesicles, and epididymis, TagBFP fluorescence was more restricted and was particularly prominent in the mucosal folds of the seminal vesicles (**Figure 6D–D′**). In adult tissues, reporter fluorescence displayed regional variation along the epididymis. Signal was detected in the caput region, whereas little or no fluorescence was readily detected in the corpus and cauda regions (**Figure 6E, E′**). TagBFP fluorescence was not readily detected in the testicular parenchyma at the stages examined.

Together, these observations indicate that *Aldh1a3* reporter expression becomes progressively restricted to selected epithelial domains of the male reproductive tract during development, including the seminal vesicles, rete testis, and caput epididymis.

*Female reproductive tract*. We next examined *Aldh1a3*-TagBFP fluorescence during female urogenital development (**Figure 7**). At E16.5, reporter fluorescence was detected along the developing uterine horns and at their caudal midline convergence dorsal to the bladder (**Figure 7A**). In contrast, TagBFP fluorescence was not readily detected in the developing ovary (**Figure 7B**). Strong fluorescence in the cortical collecting ducts of the embryonic kidney provided an internal positive reference for reporter detection in the same samples (**Figure 7B**).

**Figure 7.**
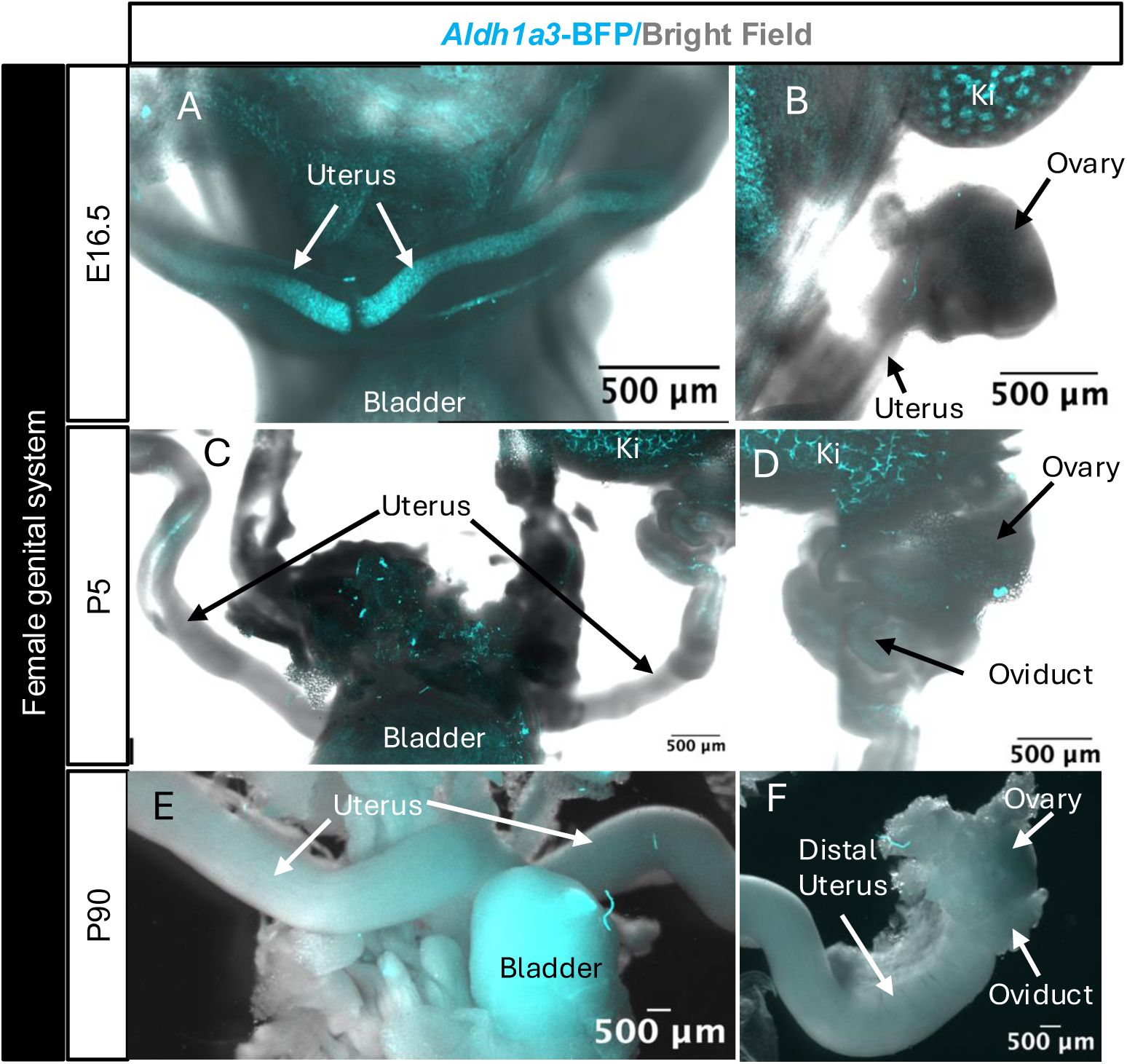
*Aldh1a3-P2A-TagBFP* reporter expression in the female reproductive tract. (**A–F**) Confocal whole-mount fluorescence images overlayed with brightfield showing *Aldh1a3*-TagBFP expression (cyan) across developmental stages of the female genital system: E16.5 embryonic stage (**A, B**), P5 postnatal stage (**C, D**), and P90 adult stage (**E, F**). Ki: Kidney. Scale bars: 500 µm.

As development progressed, TagBFP fluorescence in the female reproductive tract appeared to decrease postnatally (**Figure 7C–F**). At adult stages, reporter fluorescence remained readily detectable in the bladder, whereas signal was not readily detected in the ovary or oviduct under the imaging conditions used (**Figure 7D–F**). These observations are consistent with a temporally and spatially restricted pattern of *Aldh1a3* reporter expression in the female reproductive tract, with detectable signal in the developing uterine horns and bladder but limited or undetectable signal in the ovary and oviduct.

## Discussion

In this study, we generated and characterized a bicistronic *Aldh1a3-P2A-TagBFP* knock-in mouse line for the direct visualization of *Aldh1a3* expression during development. The *TagBFP* cassette was inserted immediately upstream of the endogenous *Aldh1a3* stop codon, placing reporter expression under the control of endogenous *Aldh1a3* regulatory elements. Targeted integration was validated by PCR and Oxford Nanopore long-read sequencing at the analyzed regions of the targeted locus. The detection of TagBFP fluorescence in multiple previously described *Aldh1a3* expression domains, together with its overlap with endogenous Aldh1a3 immunoreactivity, supports the use of this allele as a fluorescent reporter of *Aldh1a3* expression during development.

### A fluorescent resource for studying *Aldh1a3* expression

Existing genetic tools for studying *Aldh1a3*-expressing cells include inducible CreERT2-based alleles and lacZ reporter models^10,11^. These approaches have provided important information about *Aldh1a3* expression patterns and, in the case of Cre-based strategies, lineage contribution. However, each approach has specific experimental considerations. CreERT2-based strategies require breeding with Cre-dependent reporter strains and temporal activation by tamoxifen, whereas lacZ-based reporters require tissue fixation and processing for detection, limiting their use for direct visualization of reporter expression in fresh tissues.

The *Aldh1a3-P2A-TagBFP* allele provides a complementary approach by enabling direct fluorescence visualization of reporter expression from the endogenous *Aldh1a3* locus in fresh tissues. Because TagBFP is spectrally distinct from commonly used green, red, and far-red fluorescent reporters, this line can also be combined with additional fluorescent markers to compare expression domains within developing tissues. This application was illustrated by combining the reporter allele with *Pax2-GFP* to visualize the spatial relationship between *Aldh1a3* and *Pax2* expression domains during urogenital development.

### *Aldh1a3* expression during kidney development

Our analysis provides a spatial characterization of *Aldh1a3* reporter expression during mesonephric and metanephric kidney development. At early developmental stages, TagBFP fluorescence was detected in the caudal nephric duct (ND) and subsequently in ureteric bud (UB)-derived structures, consistent with previously reported *Aldh1a3* expression domains^1,4,9^.

Previous studies of mesonephric and ND development have largely relied on broad lineage markers, including *Pax2* and *Hoxb7*^19–22^, which label the entire ND epithelium and its derivatives but do not resolve the molecular heterogeneity or dynamic behavior of distinct ND progenitor populations^9^. Our previous single-cell transcriptomic analysis identified *Aldh1a3* as a marker of the NdPr4 ND progenitor population at E9.5^9^. The present reporter line provides complementary anatomical information by enabling visualization of the corresponding *Aldh1a3*-positive domain in intact embryos. The presence of TagBFP fluorescence in the caudal ND and in cells located at the leading edge of ND migration is consistent with the previously described localization of the NdPr4 population^9^. The *Aldh1a3-P2A-TagBFP* reporter line provides a unique opportunity to directly visualize and characterize this *Aldh1a3*-expressing population, and in combination with *Pax2*-GFP reporter enabling investigation of NdPr4 emergence, maturation, and cellular behaviors during ND development.

At later developmental stages, reporter fluorescence was detected in UB-derived structures, including components of the branching collecting duct system. The distribution of fluorescence was heterogeneous and appeared enriched in selected regions of the developing kidney. In contrast, TagBFP fluorescence was not readily detected in neighboring nephron progenitor compartments under the imaging conditions used. These observations are consistent with preferential *Aldh1a3* expression in UB-derived epithelial populations^4,14^. The reporter also revealed TagBFP fluorescence in additional mesonephric structures at E12.5, including the anterior ND, mesonephric tubules, and developing Müllerian duct. These observations extend the anatomical description of *Aldh1a3* expression during urogenital development.

Retinoic acid (RA) signaling plays important roles in kidney and urogenital development, including ureteric bud formation and branching morphogenesis, and genetic studies have demonstrated functional overlap between *Aldh1a3* and *Aldh1a2* during kidney development^4,15^. The *Aldh1a3-P2A-TagBFP* allele provides a spatial framework for examining *Aldh1a3* expression across the developing urogenital system.

### *Aldh1a3* expression in the reproductive tract

The reporter line also revealed dynamic and tissue-specific *Aldh1a3* reporter expression patterns within the developing reproductive system. In males, TagBFP fluorescence was detected in several urogenital structures, including the rete testis, seminal vesicles, and ductus deferens, consistent with the developmental relationships between the ND and its reproductive tract derivatives^1,4,23^. Postnatally, reporter fluorescence was particularly prominent in the seminal vesicle epithelium and displayed regional variation along the epididymis.

While *Aldh1a3* expression in the seminal vesicle epithelium is consistent with previous reports in the postnatal male genital tract^1^, our reporter provides additional spatial resolution of *Aldh1a3* expression in the developing male reproductive tract, including the caput epididymis and rete testis. Notably, we detected reporter fluorescence in the rete testis as early as E12.5, extending previous transcriptomic and immunohistochemical evidence of *Aldh1a3* expression in the rete testis at E14.5^24,25^. These expression domains provide a resource for investigating the potential roles of Aldh1a3 and RA signaling in epithelial differentiation and regional specialization during reproductive tract development.

In females, TagBFP fluorescence was detected in the developing uterine horns, including at E16.5, and in the bladder, whereas reporter fluorescence was not readily detected in the ovary or oviduct under the conditions examined. These findings indicate that *Aldh1a3* reporter expression is spatially restricted within the developing female urogenital system. RA signaling is known to contribute to Müllerian duct and uterine development, with previous studies highlighting a prominent role for stromal Aldh1a2-mediated RA production^16,17^. Our detection of *Aldh1a3* reporter fluorescence in the developing uterine horns provides additional spatial information on *Aldh1a3* expression during female reproductive tract development.

### Limitations and future applications

Several limitations should be considered when interpreting this reporter line. TagBFP fluorescence reflects reporter expression from the *Aldh1a3* locus and may not fully reproduce endogenous Aldh1a3 protein abundance, localization, or enzymatic activity. The P2A strategy is expected to generate separate Aldh1a3 and TagBFP proteins from the same bicistronic transcript; however, differences in protein stability, translation efficiency, or detection sensitivity may influence reporter output. In addition, fluorescence detection depends on reporter expression levels, tissue properties, and imaging conditions, such that weak *Aldh1a3* expression domains may be less readily detected. Therefore, the allele should be interpreted primarily as a reporter of *Aldh1a3* expression rather than as a direct readout of Aldh1a3 enzymatic activity or RA signaling, and absence of detectable fluorescence should not be interpreted as definitive evidence of absent *Aldh1a3* expression. The present study does not directly assess RA levels or downstream RA signaling.

Unlike Cre-based approaches, this reporter provides a direct fluorescent readout from the endogenous *Aldh1a3* locus without requiring additional reporter crosses or tamoxifen-mediated recombination. However, this allele does not provide permanent lineage labeling, and lineage-tracing experiments will require complementary genetic strategies^10,11^. With appropriate validation, TagBFP fluorescence could be used to identify and isolate *Aldh1a3*-expressing populations for downstream molecular and cellular analyses.

Overall, this study establishes and characterizes a fluorescent *Aldh1a3* knock-in reporter mouse and provides a detailed map of *Aldh1a3* reporter expression across developing kidney and reproductive tissues. The *Aldh1a3-P2A-TagBFP* allele enables direct visualization of *Aldh1a3* expression in developing and adult tissues and provides a complementary tool for studying its spatial distribution.

## Materials and Methods

### Animals

All animal experiments performed in this study were conducted in compliance with the Canadian Council of Animal Care ethical guidelines and were approved by the Université de Sherbrooke and McGill Animal Care Committee. Mice were housed in autoclaved cages with free access to food and water, as well as appropriate and sufficient nesting and bedding material. Mice had a 14-hour cycle of light and darkness. Mouse rooms and cages were well ventilated and kept at a temperature range of 20-24°C, with a relative humidity of 45-65%.

### Gene Targeting and Generation of knocking mice by CRISPR/Cas9 technology

#### Knock-in donor vector construction and gRNA design

A 1,943-base pair DNA fragment containing the 5′ homology arm upstream of the target locus, the CRISPR target site, a *P2A* sequence, the *TagBFP* coding region, an *SV40 poly(A)* signal, and the 3′ homology arm downstream of the *Aldh1a3* stop codon (**Figure 1A**) was commercially synthesized and cloned into the pUC57 vector at the *EcoRV* site by GeneScript.

Guide RNA (gRNA) target sequences were designed using the CRISPOR online tool to identify candidate target sites located as close as possible to the desired insertion site within the *Aldh1a3* locus. Candidate gRNAs were selected based on high predicted on-target activity and minimal off-target potential, prioritizing guides with at least two nucleotide mismatches to predicted off-target genomic sites. Guide RNA activity was experimentally validated by microinjection of CRISPR/Cas9 reagents into mouse zygotes, followed by embryo culture to the blastocyst stage. Genomic DNA was isolated from individual blastocysts, and editing efficiency was assessed by PCR amplification of the target locus followed by Sanger sequencing. The gRNA exhibiting the highest editing efficiency was selected for generation of the *Aldh1a3-P2A-TagBFP* knock-in mouse line.

#### Microinjection

All embryos were obtained from superovulated WT C57BL/6 female mice (Charles River) following standard hormonal stimulation with pregnant mare serum gonadotropin (PMSG) and human chorionic gonadotropin (hCG). Embryos were generated by in vitro fertilization (IVF). Oocytes were collected at embryonic day 0.5 (E0.5) and fertilized by IVF and cultured overnight in Advanced KSOM medium (Sigma) under standard conditions. At the 2-cell stage, genome editing reagents were delivered by nuclear microinjection into both blastomeres. Each nucleus was injected with a mixture containing 50 ng/µL Cas9, 50 ng/µL guide RNA (gRNA), and 20 ng/µL donor DNA. Following microinjection, embryos were cultured to assess viability and subsequently transferred into the oviducts of pseudopregnant CD1 recipient females for further development.

### Genomic PCR and sequencing analysis

#### Mouse model generation and PCR-based genotyping strategy

Founder mice were generated on a C57BL/6 background and identified by PCR-based genotyping followed by Sanger sequencing. Genomic DNA was extracted from tail biopsies using standard proteinase K digestion and ethanol precipitation and used as a template for PCR amplification. Primers were designed to amplify external and internal junctions of the targeted locus (**Table S1**). Specifically, four diagnostic junctions were analyzed to confirm correct targeted integration and allele integrity: the 5′ and 3′ external junctions, and the 5′ and 3′ internal junctions. PCR products were resolved by agarose gel electrophoresis.

To assess sgRNA efficiency, transient blastocysts generated by RNP-only injections (omitting the DNA donor plasmid) were genotyped by PCR amplifying extracted genomic DNA using primers F4/R4 (**Table S1**), followed by Sanger sequencing. Targeting efficiency and indel frequencies were assessed using the Tracking of Indels by Decomposition (TIDE) web tool.

All mice were maintained on a C57BL/6 genetic background and routinely genotyped using the F2-R2 primer pair (**Table S1**) to discriminate wild-type, heterozygous, and homozygous animals.

#### Oxford Nanopore long-read sequencing validation of the Aldh1a3-P2A-TagBFP allele

*Aldh1a3-P2A-TagBFP* homozygous mice were validated by Oxford Nanopore long-read sequencing using amplicons generated with the same external and internal junction primer sets described above. Long-read sequencing was performed to confirm correct integration of the *P2A-TagBFP-Poly(A)* cassette, evaluate the structural integrity of the targeted allele, and exclude potential complex rearrangements at the insertion site.

A total of 400 ng of purified genomic DNA was used to prepare sequencing libraries using the Rapid Barcoding Kit (SQK-RBK114.94, Oxford Nanopore Technologies) according to the manufacturer’s instructions. The final libraries were loaded onto a PromethION flow cell for long-read sequencing. Raw sequencing data were basecalled using the Super Accuracy (SUP) model in Dorado. The resulting FASTQ files were analyzed using a bioinformatics pipeline developed by the University of Sherbrooke RNomics platform. Sequencing reads were aligned to the reference genome, and the targeted locus was visualized using Integrative Genomics Viewer (IGV) for manual inspection of the insertion site, junction regions, and overall structural integrity of the targeted allele.

### Tissue Preparation and Fluorescence Microscopy

*Aldh1a3-P2A-TagBFP* and *Pax2-GFP* transgenic mouse embryos on a C57BL/6 background were obtained via natural mating, with noon on the day of vaginal plug detection designated as embryonic day 0.5 (E0.5). Whole embryos and whole-mount urogenital systems were dissected in phosphate-buffered saline (PBS), and kidneys were sectioned at 200 μm using a Leica VT1200S vibratome. Fluorescence images were acquired using a SteREO Lumar.V20 stereomicroscope (Carl Zeiss) and an FV3000 confocal laser scanning microscope (Olympus) equipped with 405 nm (for TagBFP) and 488 nm (for GFP) laser lines.

### Whole-mount immunofluorescence staining

Whole mount immunofluorescence staining of *Aldh1a3-P2A-TagBFP* mouse embryos was performed using the protocol described in ^9,26^. Primary antibodies and dilutions used are as follows: Aldh1a3 (1:100, Millipore Sigma, ABN427); anti-mTagBFP (1:100, Antibodies online, ABIN7272956). Whole mount imaging of *Aldh1a3-P2A-TagBFP* mouse embryos was performed on a SteREO Lumar V20 Carl Zeiss stereomicroscope. Images were analysed with Image J (Fiji) 2.16.0/1.54p software version. Three-dimensional rendering of confocal image stacks from the E11.5 *Aldh1a3-P2A-TagBFP;Pax2-GFP* urogenital system was performed using arivis Vision4D software (ZEISS).

### Statistical analysis

Genotype distributions were compared with expected Mendelian inheritance ratios using a chi-square goodness-of-fit test. A p value < 0.05 was considered statistically significant.

## Acknowledgements

We express our sincere gratitude to Dr. Yojiro Yamanaka for facilitating the financial and administrative maintenance of the mouse colony during the transitional phase while O.S.-F. established his independent laboratory at the Université de Sherbrooke. We thank Daniel Garneau for his exceptional technical support and training in fluorescence microscopy and image analysis. We thank Dr. Rodolphe Soret for his constructive suggestions throughout this study. We also thank all members of the Sanchez-Ferras laboratory for helpful discussions and specially Sarah-Gabrielle Taillander Pensarini for colony management. We are grateful to the McGill University Transgenic Core Facility (Mitra Cowan, Nobuko Yamanaka, Hedyeh Rahimian) for expert technical assistance in generating the mouse line, as well as the RNomics platforms at the Université de Sherbrooke for technical support.

## Author contributions

A.C. performed research; A.R.O. performed research; L.B. performed research; E.C.-M. analyzed data; M.H. performed research. M.B. supported the initiation of the project and provided funding that supported the initial generation and rederivation of the reporter line. O.S.-F. conceived and directed the research, performed research, analyzed data. The manuscript was written by O.S.-F and edited by A.C., A.R.O., L.B., E.C.-M and M.H.

