## Supplementary information for "A bicistronic *Aldh1a3-P2A-TagBFP* knock-in reporter mouse line for studying genitourinary tract development"

### Supplementary Figures

| Target Name | Spacer Sequence (5'-3') | PAM |
|---|---|---|
| *Aldh1a3* gRNA | CTCGAGGAGAAGAACCCCTG | AGG |

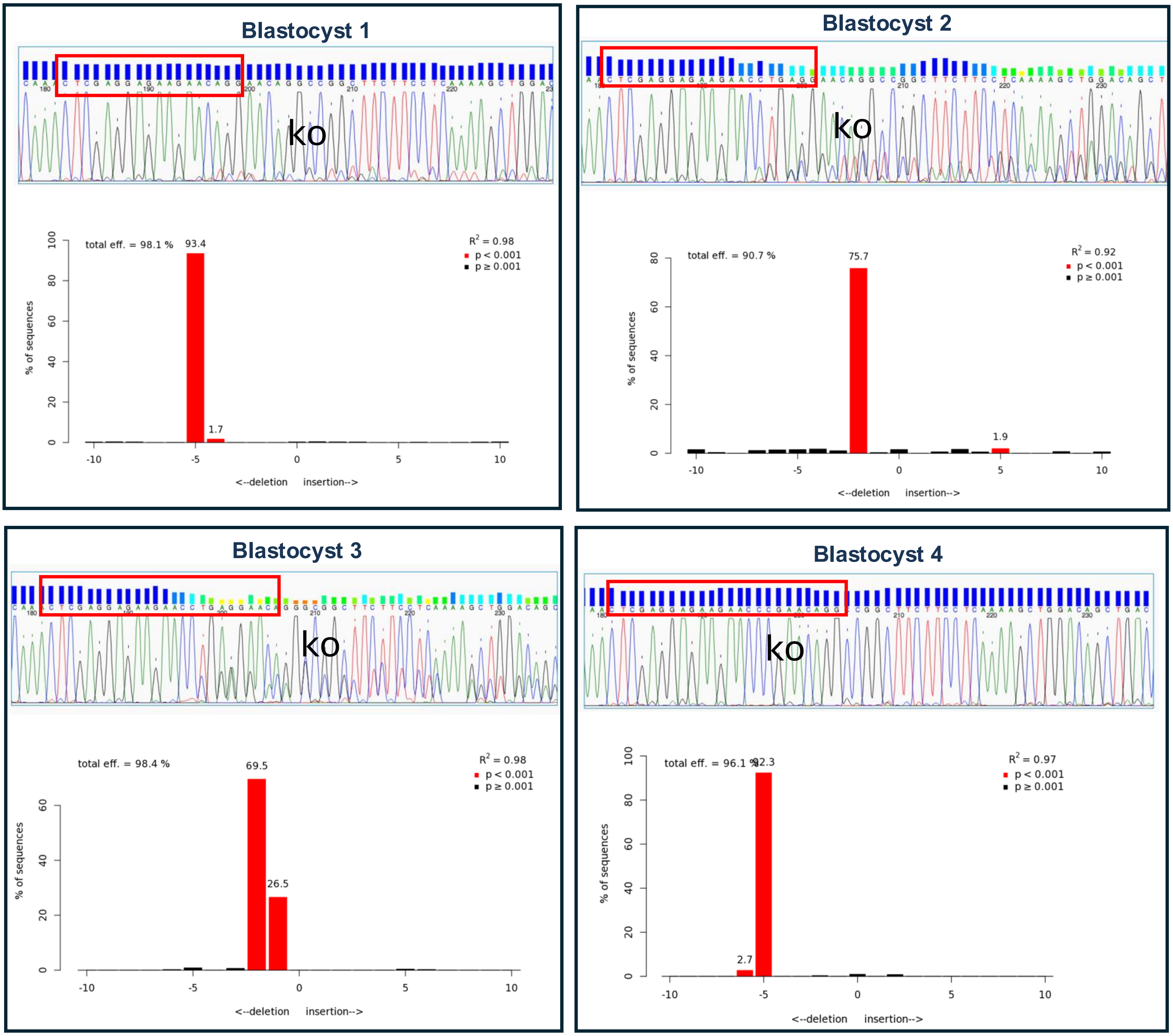


**Figure S1. Validation of *Aldh1a3* sgRNA targeting efficiency in transient blastocysts.** (Top Table) Target name, spacer sequence, and protospacer adjacent motif (PAM) for the designed *Aldh1a3* single-guide RNA (sgRNA). (Main Panels) Sanger sequencing chromatograms of PCR products generated with primers F4-R4 and corresponding Tracking of Indels by Decomposition (TIDE) analysis plots for four representative transient blastocysts (Blastocysts 1–4). Red boxes on the chromatograms outline the sgRNA target sequence. Histograms illustrate the frequency and spectrum of insertions and deletions (indels) relative to the predicted cleavage site, with total editing efficiency percentages shown and statistically significant indels ($p < 0.001$) highlighted in red.

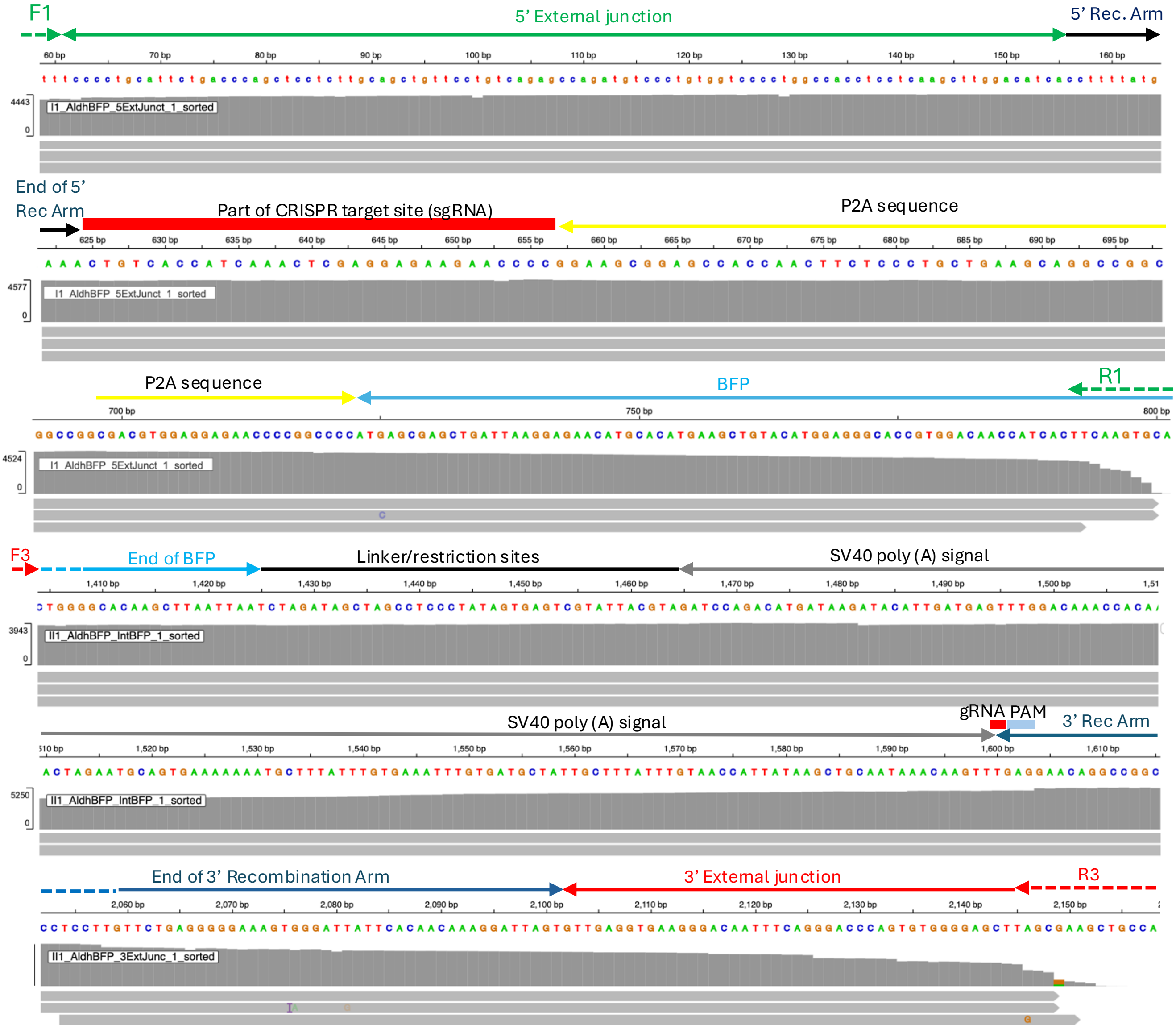


**Figure S2. Oxford Nanopore long-read sequencing validation of the *Aldh1a3-P2A-TagBFP* knock-in allele.** Integrative Genomics Viewer (IGV) alignment map of Oxford Nanopore long reads mapped across the targeted *Aldh1a3* locus to confirm precise integration and allele integrity. Annotations above the nucleotide sequence highlight key genetic elements: the 5’ external junction (green arrow), 5' homology/recombination arm (black arrow), CRISPR target site/sgRNA sequence (red box), *P2A* self-cleaving peptide sequence (yellow arrow), *BFP* reporter sequence (light blue arrow), linker and restriction enzyme sites (black line), *SV40* polyadenylation signal (grey arrow), gRNA PAM site (red and blue box), 3' homology/recombination arm (dark blue arrow), and 3’ external junction (red arrow). Primers used for amplification are indicated as F1, R1, F3, and R3. Coverage tracks (grey histograms) demonstrate continuous, full-length sequence alignment across all internal and external junctions.

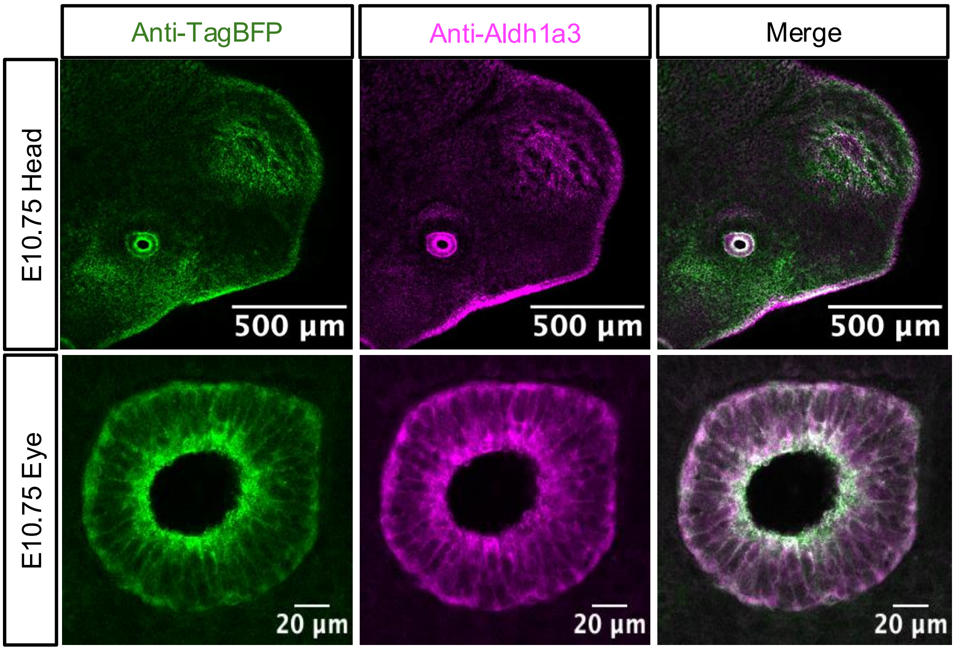


**Figure S3. Co-localization of TagBFP reporter and endogenous Aldh1a3 protein.** Whole-mount immunostaining of E10.75 embryos displaying anti-TagBFP signal (green; left), anti-Aldh1a3 protein signal (magenta; middle), and merged channel overlays (right) in the cranial region (**top row**) and high-magnification views of the developing eye (**bottom row**). Scale bars: 500 µm (top), 20 µm (bottom).

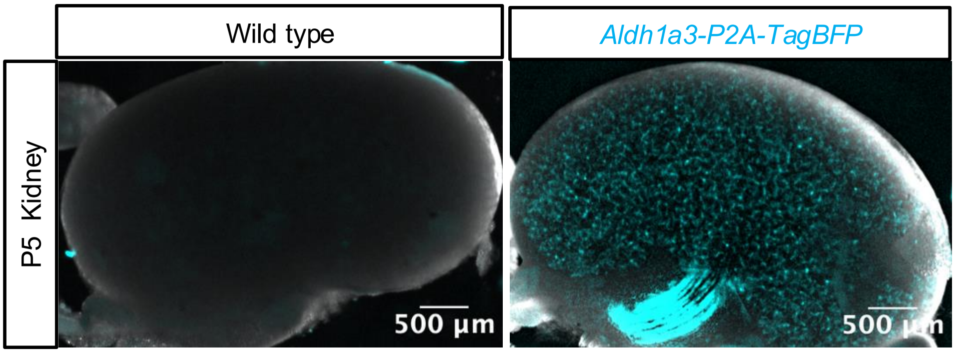


**Figure S4. Specificity of *Aldh1a3-P2A-TagBFP* reporter fluorescence in postnatal kidneys.** Representative fluorescence and brightfield overlay images of wild-type controls (left) and *Aldh1a3-P2A-TagBFP* reporter P5 kidneys imaged using a V20 stereomicroscope. Control and reporter kidneys were imaged using identical acquisition settings within each imaging modality. Scale bars: 500 µm.

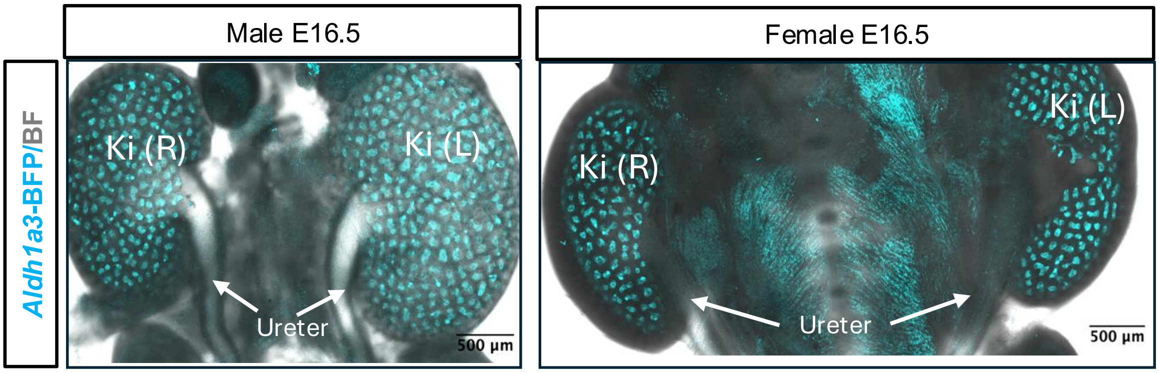


**Figure S5. *Aldh1a3-P2A-TagBFP* reporter expression in E16.5 male and female kidneys.** Whole-mount fluorescence images overlayed with brightfield showing *Aldh1a3*-TagBFP expression (cyan) in right and left kidneys isolated from E16.5 male (left panel) and female (right panel) *Aldh1a3-P2A-TagBFP* homozygous embryos. Labels indicate the left and right kidneys and ureters. Ki (R): Right kidney; Ki (L): Left kidney. Scale bars: 500 µm.

### Table S1. Primers used

| Primer name | Primer Sequence (5'-3') |
|---|---|
| F1 | 5’-TGGGCTACTATCTGCTCCCTT-3’ |
| R1 | 5’-CCTCGGATGTGCACTTGAA-3’ |
| F2 | 5’-CATCGTAGAGCCTACTGCCTT-3’ |
| R2 | 5’-CCCATATCCTATCCGTCTGCC-3’ |
| F3 | 5’-ATACTGCGACCTCCCTAGCA-3’ |
| R3 | 5’-TTTCCATGGCAGCTTCGCTA-3’ |
| F4 | 5’-TCACACGCAGGAACTTAGCAT-3’ |
| R4 | 5’-TCCTATCCGTCTGCCCACA-3’ |
